# Novel directly and co-operatively drive lytic polysaccharide monooxygenase activity with AA8 module

**DOI:** 10.1101/2025.09.02.673750

**Authors:** Yuxing Sun, Congcong Yuan, Liangkun Long, Shaojun Ding

## Abstract

The single-domain auxiliary activity family 12 (AA12) pyrroloquinoline quinone-dependent oxidoreductases and free AA8 modules are prevalent in cellulolytic fungi, however, their function in polysaccharide biodegradation is still confused. Here, we characterized three single-domain AA12 oxidoreductases and one free AA8 module from *Thermothelomyces thermophilus* and *Thermothielavioides terrestris*. All three single-domain AA12 oxidoreductases are restrict dehydrogenases with trace oxidase activity. All three single-domain AA12 enzymes could directly transfer electrons to lytic polysaccharide monooxygenase (LPMO) and drive *Nc*LPMO9C activity. Furthermore, inter-protein electron transfer between single-domain AA12 enzymes and the AA8 module was observed. The AA12 enzyme-driven *Nc*LPMO9C efficiency could be significantly enhanced by the addition of free AA8 module *Tth*AA8B, probably attributing to the acceleration of electron transfer from AA12 enzymes to *Nc*LPMO9C and the attenuation of H_2_O_2_ accumulation mediated by *Tth*AA8B. Our findings highlight the potential role of single-domain AA12 enzyme and free AA8 modules in the biodegradation system of LPMOs.

**IMPORTANCE:** This study reveals the directly electron transferring and driving capability of single-domain AA12 PQQ-dependent enzyme for the oxidative reaction of lytic polysaccharide monooxygenase (LPMO). This funding is quite distinct from the AA8 cytochrome domain-dependent driving pattern of the previous characterized multi-domain *Cc*PDH. We also demonstrated that this priming capability could be facilitated by the free AA8 cytochrome module, providing new insight on the interactions and functions of single-domain AA12 enzymes and free AA8 modules in fueling LPMO activity during fungal lignocellulose biodegradation process. These findings collectively provide evidence for the potential function of widespread single-domain AA12 PQQ-dependent enzymes and free AA8 cytochrome modules as unique enzyme redox partners in cellulolytic fungi.

## INTRODUCTION

The first fungal pyrroloquinoline quinone (PQQ)-dependent enzyme, now classified as an Auxiliary Activity Family (AA12) pyranose dehydrogenase (*Cc*PDH), was discovered in the basidiomycete fungus *Coprinopsis cinerea* in 2014 (1). It primarily catalyzes the oxidation of 2-keto-D-glucose (2KG), L-fucose, and several rare sugars—including D-arabinose, L-galactose, D-talose, and L-gulose. In contrast, it shows no activity toward common sugars such as glucose or cellobiose (2).

*Cc*PDH consists of an N-terminal AA8 cytochrome b domain, the catalytic PQQ domain, and a C-terminal cellulose-binding domain connected by two flexible linker peptides (1). The three-domain structure of *Cc*PDH resembles certain fungal cellobiose dehydrogenases (CDHs), in which the AA8 cytochrome b domain was linked to an AA 3 flavin-containing sugar dehydrogenase domain (3–6).

CDH plays a pivotal role in lytic polysaccharide monooxygenase (LPMO)-mediated lignocellulose degradation due to the efficient inter-domain electron transfer capability of AA8 domain (7–9). The spectral and electrochemical properties of the AA8 cytochrome b domain in *Cc*PDH are almost identical to those found in CDHs (2–4). Similarly, the inter-domain electron transfer from the PQQ cofactor in the catalytic domain to the heme b in the cytochrome domain occurs in *Cc*PDH, thereby driving *Neurospora crassa Nc*LPMO9C and *Nc*LPMO9F action (10, 11). Thus, PQQ-dependent oxidoreductases have attracted considerable interest in recent years due to their proposed role in plant cell wall degradation by acting as critical auxiliary enzymes for cellulolytic enzymes (5, 12). However, previous study revealed that this activation depends strongly on AA8 domain, since the truncation of AA8 domain almost completely abolished the LPMO activity and AA12 enzyme activity, indicating that the AA8 domain is needed for mediating electron transfer from the PQQ cofactor in the AA12 domain to the LPMO (11).

In nature, AA12 PQQ-dependent oxidoreductases predominantly exist as a single-domain enzyme, with a minority linked to AA8 domains via a flexible linker peptide (https://www.cazy.org/AA12.html) (13). Notably, AA8 modules can also exist as an independent module (https://www.cazy.org/AA8.html) (13). So far, only two AA12 PQQ-dependent oxidoreductases, including *Cc*PDH from *C. cinerea* and *Tr*AA12 from *Trichoderma reesei* have been characterized (1, 14). Despite the multi-modular *Cc*PDH from *C. cinerea* was extensively studied and its capability of activating LPMO activity was confirmed, the functional roles of majority of single-domain fungal AA12 PQQ-dependent oxidoreductases and its interplay with single AA8 module remain poorly understood. Recently, Momeni et al. discovered a novel AA7 cellooligosaccharide dehydrogenase *Fg*CelDH7C from *Fusarium graminearum*, containing flavin-adenine dinucleotide (FAD)-binding domain but without AA8 domain. Interestingly, unlike other known sugar dehydrogenases that depend on an additional cytochrome haem domain to prime LPMOs, AA7 *Fg*CelDH7C can directly transfer electrons to *Podospora anserina Pa*LPMO9E and *Pa*LPMO9H, thereby fueling cellulose degradation without the need for exogenous reductants (15). This finding not only expands our understanding of the functional diversity within AA7 proteins but also sheds new light on the functionality of the single-domain AA12 PQQ-dependent oxidoreductases in fungi.

In this study, three single-domain AA12 PQQ-dependent oxidoreductases and two free AA8 modules were heterologously expressed in *Pichai pastoris*. We hypothesized that the single-domain AA12 enzymes could directly or co-operatively with AA8 module transfer electrons to LPMO and drives LPMO activity. To this end, the redox states of AA8 modules were monitored by spectral scanning and stopped-flow experiment, while electron transfer between AA12 enzymes and LPMO was analyzed using electron paramagnetic resonance (EPR). Additionally, the synergy of AA12 enzymes or/and AA8 module with C4-oxidizing *Nc*LPMO9C in degrading phosphoric acid-swollen cellulose (PASC) was investigated. This study demonstrated that single-domain AA12 PQQ-dependent oxidoreductases alone can transfer electron to *Nc*LPMO9C and activate *Nc*LPMO9C action. Furthermore, the presence of free AA8 module significantly enhanced the oxidative activity of LPMO in AA12 enzyme-AA8 module-*Nc*LPMO9C system compared to AA12 enzyme-*Nc*LPMO9C system, suggesting the inter-molecular interplay between the single-domain AA12 PQQ-dependent dehydrogenase, free AA8 module and LPMO.

## MATERIALS AND METHODS

### Materials

The *P. pastoris* secretory expression vector pPICZαA was purchased from Invitrogen (Carlsbad, CA, USA). *Escherichia coli* Top10 competent cells (for plasmid amplification) and *Pichia pastoris* X-33 strain (eukaryotic expression host) were obtained from TransGen Biotech (Beijing, China). Lytic polysaccharide monooxygenase (*Nc*LPMO9C) was prepared in-house as previously described (16). Catalase (CAT) and horseradish peroxidase (HRP) were sourced from Aladdin Biochemical Technology (Shanghai, China). Restriction enzyme *Sac* Ⅰ and plasmid extraction kits were purchased from Takara Bio (Tokya, Japan). PCR purification kits and BCA protein assay kits were acquired from TransGen Biotech (Beijing, China). 2,6-Dichloroindophenol (DCIP), PQQ, and L-fucose were obtained from Aladdin (Shanghai, China). Zeocin was supplied by Invitrogen. Yeast extract, peptone, and tryptone were procured from Oxoid (Basingstoke, UK). 2,6-Dimethoxyphenol (2,6-DMP), Amplex Red, sodium dithionite, ascorbic acid (AscA), phosphoric acid, and 30% H₂O₂ were purchased from Sinopharm Chemical Reagent Co., Ltd (Beijing, China). All other chemicals were analytical-grade domestic reagents. PASC was prepared from Avicel PH-101 using the method described by Zhang et al. (17).

### Expression and purification of enzymes

The complete amino acid sequences of two AA12 PQQ-dependent oxidoreductases (GenBank No. AEO55799.1 and AEO59868.1; designed *Tth*AA12A and *Tth*AA12B) and two AA8 modules (GenBank No. AEO59519.1 and AEO61106.1; designed *Tth*AA8A and *Tth*AA8B) from *Thermothelomyces thermophilus* ATCC 42464, as well as one AA12 PQQ-dependent oxidoreductase (GenBank No. AEO64217.1; designed *Tte*AA12A) from *Thermothielavioides terrestris* NRRL 8126, were retrieved from the National Center for Biotechnology Information (NCBI) database (https://www.ncbi.nlm.nih.gov/) (Figure S1). These gene fragments were synthesized by Sangon

Biotech (Shanghai, China) with modified codons according to the codon preference of *P. pastoris*. The target DNA fragments were cloned into the pPICZαA expression vector (Invitrogen) via *Bst*BI/*Eco*RI restriction sites, replacing the native α-factor sequence with the corresponding signal peptide sequences. The recombinant plasmids were linearized with *Sac*I and transformed into *Pichia pastoris* X-33 competent cells via MicroPulser Electroporator (Bio-Rad, Hercules, CA, USA). Transformants were selected on YPDZ agar plate (containing 1% yeast extract, 2% peptone, 2% glucose, 2% agar, and 100 μg/mL Zeocin) at 28°C for 72 h. Single colonies were inoculated into BMGY medium (1% yeast extract, 2% peptone, 1% glycerol, 1.34% yeast nitrogen base, and 4×10^5^% biotin in 100 mM potassium phosphate buffer, pH 6.0) and cultured at 28°C with shaking at 250 rpm. When the OD_600_ reached 2.0, cells were transferred to BMMY induction medium (with 1% methanol replacing glycerol) for 6 days, with daily methanol supplementation to maintain 1% concentration.

Recombinant proteins with a C-terminal 6×His tag were purified by a nickel-nitrilotriacetic acid (Ni–NTA) superflow resin column (Qiagen, Valencia, CA, USA), then dialyzed at 4°C overnight using 14 kDa MWCO membranes (Viskase, Lombard, IL, USA) in 200 mM sodium phosphate buffer (pH 7.0). Reconstitution of the apoenzyme of purified *Tth*AA12A, *Tth*AA12B and *Tte*AA12A was carried out by incubating the enzymes with 0.2 mM PQQ and 2 mM CaCl_2_ at room temperature for 60 min. Excess PQQ and calcium ions were removed through three additional dialysis cycles (8 h each). All purified 6xHis-tagged enzymes were stored at −80°C until use. Protein concentration was determined using a BCA assay kit (Thermo Scientific, Waltham, MA, USA), and purity was verified by 12% SDS-PAGE.

### Phylogenetic tree, sequence alignment analysis and structure modeling

To explore the relationship of *Tth*AA12A, *Tth*AA12B and *Tte*AA12A with other AA12 PQQ-dependent oxidoreductases, the amino acid sequences of 120 single-domain AA12 enzymes along with two characterized AA12 enzymes were retrieved from the GenBank database in the NCBI. The phylogenetic tree was constructed using the neighbor-joining algorithm in MEGA 7.0 (8), with subsequent visualization refinement using iTOL (https://itol.embl.de/) (18, 19). For structural characterization, three-dimensional models of both AA12 enzymes and AA8 modules were constructed by SWISS-MODEL. Structural visualization and comparative analyses were performed using PyMOL Molecular Graphics System (version 2.1.0, Schrödinger, LLC). Multiple amino acid sequence alignment of the three AA12, *Tr*AA12, and *Cc*PDH) was carried out using ESPript 3.0 (https://espript.ibcp.fr/ESPript/ESPript/index.php) to identify conserved motifs and sequence variations (20).

### Enzymatic activity

The activity of AA12 enzymes was determined in a reaction mixture (1 mL) containing 100 mM L-fucose, 200 μM DCIP, and 0.1 μM AA12 enzymes in 50 mM sodium acetate buffer (pH 6.0). Reactions were conducted at 50°C for 4 min and terminated by adding 100 mM sodium dodecyl sulfate (SDS). DCIP reduction was monitored spectrophotometrically at 530 nm. One unit of enzyme activity (IU) was defined as the amount of enzyme required to reduce 1 μmol of DCIP per minute under the specified condition. The AA8 module’ activity was evaluated by spectral analysis. Absorption spectra of 10 μM AA8 module were recorded before and after reduction with sodium dithionite using X-6 ultraviolet-visible spectrophotometer (METASH, Shanghai, China) covering the wavelength range of 380–600 nm with 1 nm resolution. All spectral measurements were performed in 100 mM 3-(N-morpholino) propanesulfonic acid (MOPS) buffer (pH 6.5) at 25°C.

### Analysis of PQQ proportion in AA12 enzymes

The molar extinction coefficient of the mature AA12 enzymes at 280 nm were calculated from their amino acid compositions using ProtParam (http://web.expasy.org/protparam/), and used for the determination of the protein concentration of the purified AA12 enzymes. PQQ occupancy in AA12 enzymes was determined by using the TCA method (21). This method involved the addition of 5% trichloroacetic acid in 60 µM enzyme to release PQQ cofactor from AA12 enzymes and spectrophotometrically measured the non-covalently bound PQQ cofactor (330 nm) in the supernatant. The amount of PQQ was calculated using the molar extinction coefficient for free PQQ (ε_330_ =5.755 mM^−1^cm^−1^). The molar PQQ loading was calculated on the basis of the concentrations of AA12 enzyme determined by the absorbance at 280 nm.

### Oxidase activity assay and hydrogen peroxide detection

The oxidase activities of AA12 enzymes were assessed by quantifying their hydrogen peroxide (H₂O₂) production rates using the HRP/Amplex Red assay method (22, 23). The reaction mixture (3 mL) consisted of 100 mM L-fucose, 100 µM Amplex Red 0.1 µM, 15 units of horseradish peroxidase (HRP), and 0.1 µM AA12 enzyme in 50 mM MOPS buffer (pH 6.0). Reactions were initiated by adding the AA12 enzyme in the reaction mixture in a standard glass cuvette. H₂O₂ production was continuously monitored by tracking the formation of resorufin at 563 nm over 20 minutes at 45 °C using an X-6 ultraviolet-visible spectrophotometer. Control reactions, in which ultrapure water or third-dialysate was substituted for the AA12 enzymes, were conducted in parallel to account for non-enzymatic H₂O₂ formation. All assays were performed in triplicate. A standard curve was constructed using H₂O₂ solutions (0–100 µM) prepared in 50 mM sodium acetate buffer (pH 5.0) containing 100 mM L-fucose. H₂O₂ concentrations in experimental samples were determined based on this calibration curve.

### Effects of pH and temperature on AA12 enzyme activity and stability

The effect of pH on AA12 enzyme activity was assayed using L-fucose as substrate with pH ranging from 4.0-9.0 in three buffer systems: 50 mM sodium acetate (pH 4.0-6.0), 50 mM MOPS (pH 6.0-8.0), and 0.2 M Tris-HCl (pH 8.0-9.0) at 40°C. Relative activities were calculated as percentages of maximum activity (set as 100%). The effect of pH on AA12 enzyme stability was assayed by incubating enzymes in 0.1M universal buffer (pH 4-10) consisting of 0.1M H_2_PO_4_,0.1M CH_3_COOH and 0.1M H_2_BO_3_ at 4℃ for different time. Residual activities were measured under standard conditions (pH 6.0, 40°C, 4 min), with untreated enzyme activity as 100% control. The effect of temperature on AA12 enzyme activity was determined from 30-70°C in 50 mM sodium acetate buffer (pH 6.0), with maximum activity defined as 100%. For the thermal stability, enzyme samples were incubated at 30°C, 40°C, and 50°C for various time intervals. Remaining activities were measured under standard conditions (pH 6.0, 40°C, 4 min), with unincubated enzyme as 100%.

### Substrate specificity and kinetics of AA12 enzymes

The substrate specificity of AA12 enzymes was determined against various sugar substrates, including L-fucose, L-ribose, D-lyxose, D-xylulose, dextran, cellobiose, D-lactose, glycerol, D-sorbitol, D-mannitol, D-glucuronic acid, D-tagatose, D-trehalose, D-mannose, D-glucose, D-galactose, L-galactose, D-xylose, D-rhamnose, D-fructose, D-arabinose, maltose, L-sorbose, D-allose, D-psicose, L-arabinose, idose, 2-deoxyribose, 3,6-dideoxymannose (tyvelose), methanol, ethanol, n-butanol, tert-butanol, n-propanol, and isopropanol, with DCIP as the electron acceptor. One unit of enzyme activity was defined as the amount of enzyme required to catalyze the reduction of 1 μmol electron acceptor per minute under standard conditions. The steady-state apparent kinetics properties of AA12 enzymes towards the three active substrates (L-fucose, L-ribose, and D-lyxose) were conducted at 40°C and pH 5.0 for 2 min with substrate concentrations ranging from 0.02 to 1.5 M. The kinetic constants (*K*_m_ and *K*_cat_) were derived by fitting the data to the Michaelis-Menten equation using GraphPad Prism 5.0.

### Molecular docking

The structure of the AA12 enzymes was built by SWISS-MODEL (https://swissmodel.expasy.org) and the structure of *Cc*PDH was downloaded from PDB (https://www.rcsb.org/). The structure of L-Fucose was obtained from PubChem (http://pubchem.ncbi.nlm.nih.gov). The structural diagram of the protein was drawn by PyMOL. Using the three-dimensional (3D) structure of AA12 enzymes as a template, Schrodinger Suite was used to dock L-Fucose, and the results with the lowest binding energy were selected for analysis. Finally, Discovery Studio Visualizer was used to visualize the interaction between ligands and AA12 enzymes, and PyMOL was used to draw the active center diagram to gain a deep understanding of the characteristics and structure of intermolecular interactions.

### Interaction study between AA12 enzymes and AA8 module Stopped-Flow experiments

The reduction rate of PQQ in AA12 enzymes and heme b in AA8 module were measured using a MOS-500 circular dichroism spectrometer coupled to an SFM-3000 stopped-flow instrument (Bio-Logic, Grenoble, France). All reactions were carried out in 50 mM sodium acetate buffer at optimal reaction condition (pH 6.0, 7.0 and 6.5 for *Tth*AA12, *Tth*AA12B and *Tte*AA12A, respectively; and 60℃ for *Tth*AA12A and *Tte*AA12A, and 55℃ for *Tth*AA12B). Enzyme and substrate solutions were degassed prior to measurements to remove oxygen unless stated otherwise. The absorbance change at 330 nm was monitored over 10 s once mixing a solution of 20 μM AA12 enzymes with 500 mM L-fucose. The heme b reduction rate in AA8 module by AA12 enzymes was determined by first reacting 10 μM AA12 enzymes with 500 mM L-fucose for 5 min under the optimal reaction conditions for AA12 enzymes as described above, followed by mixing with 10 μM AA8 modules. The absorbance change at 563 nm was recorded over 40 s. Absorbance-time (Abs-t) data were fitted to an exponential function using GraphPad Prism 5.0 to obtain the reduction rates constants (*k*_obs_) of PQQ and heme b.

### Spectroscopic scanning

The absorption spectra of AA8 modules in both the oxidized state and the reduced state were monitored using a UV-Vis spectrophotometer. The enzymatic reduction of AA8 modules were conducted in 50 mM MOPS buffer (pH 6.5) at 45℃. Equimolar concentrations of AA12 enzymes and AA8 modules (10 µM each) were premixed, and the reaction was initiated by adding L-fucose to a final concentration of 100 mM. The spectral scanning was performed at 5-minutes intervals covering the wavelength range of 450–600 nm with 1 nm resolution. The fully reduced state of AA8 modules was achieved by the addition of excess sodium dithionite to the mixture according to the reference (24).

### Activation of *Nc*LPMO9C by AA12 enzymes alone or cooperatively with AA8 modules

The C4-oxidizing *Nc*LPMO9C were used to investigate the driving efficiency of AA12 enzymes or/and AA8 modules. The reaction mixture (2 mL), containing 1 μM *Nc*LPMO9C, 1 μM AA12 enzymes or/and AA8 modules with 200 mM L-Fucose, was incubated with 4 mg/mL PASC at pH 6.0 and 45°C for 12 h or 24 h with shaking at 200 rpm. The control experiments were carried out using equimolar inactivated enzymes or PQQ in same condition. Positive controls with AscA as the electron acceptor were also carried out in parallel in each AA12-driven *Nc*LPMO9C experimental group. Reactions were terminated by boiling at 99 °C for 10 min, followed by centrifugation at 10,000 rpm for 10 min. Products in the supernatant were analyzed by HPAEC-PAD as previously described, ²⁵ and the total peak area of C4-oxidized products (eluting between 18–20 min) was quantified using Chameleon 6 software (Thermo Fisher Scientific). To rule out potential effects of H₂O₂ generated by AA12 enzymes on *Nc*LPMO9C activity, 10 μg of catalase (CAT) was added to relevant reaction mixtures.

### Electron Paramagnetic Resonance (EPR)

EPR measurements were conducted using a Bruker A300-10/12 spectrometer ((Bruker) equipped with an ER4102ST standard rectangular cavity and an Oxford Instruments ESR 900 helium flow cryostat (Oxfordshire, UK). All spectra were acquired at 77 K under the following conditions: microwave frequency 9.453 GHz, power 19.73 mW, modulation amplitude 3 mT, and modulation frequency 100 kHz. Each spectrum represents an average of 4 accumulated scans to enhance the signal-to-noise ratio. AA12 enzymes and *Nc*LPMO9C were prepared in 50 mM MOPS buffer (pH 6.5) and incubated at 45 °C for 10 minutes under the following conditions: (i) *Nc*LPMO9C (100 µM) with L-fucose (200 mM); (ii) same as (i) with the addition of AA12 enzymes (10 µM). Sample preparation was performed under anaerobic conditions in a Jacomex glove box (Wuhan, China), and samples were flash-frozen in liquid nitrogen prior to measurement (15).

### Statistical analyses

Statistical significance was annotated using asterisks (*) based on p-values: **\***: p < 0.05; **\*\***: p < 0.01; **\*\*\***: p < 0.001. Excel’s =T.TEST (array1, array2,2,3) was used for two-tailed p-value calculation. Cross-validated with SPSS (Cohen’s κ=0.91).

## RESULTS

### Sequences alignment and structure modeling

Phylogenetic tree analysis of AA12 enzymes does not exhibit distinct clustering, with numerous branches that are relatively dispersed and show low similarity between clusters (Figure 1A).

**Figure 1.**
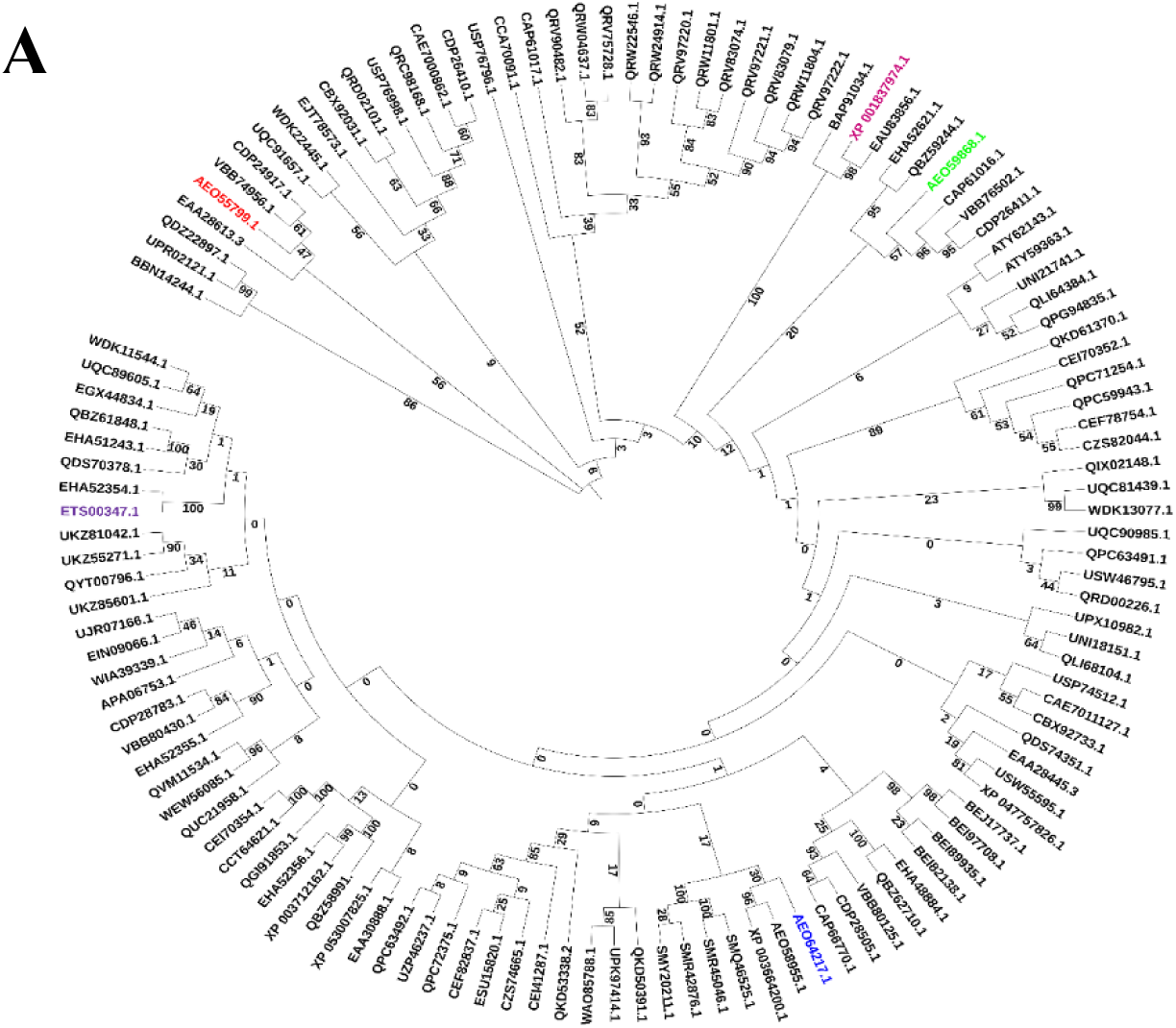
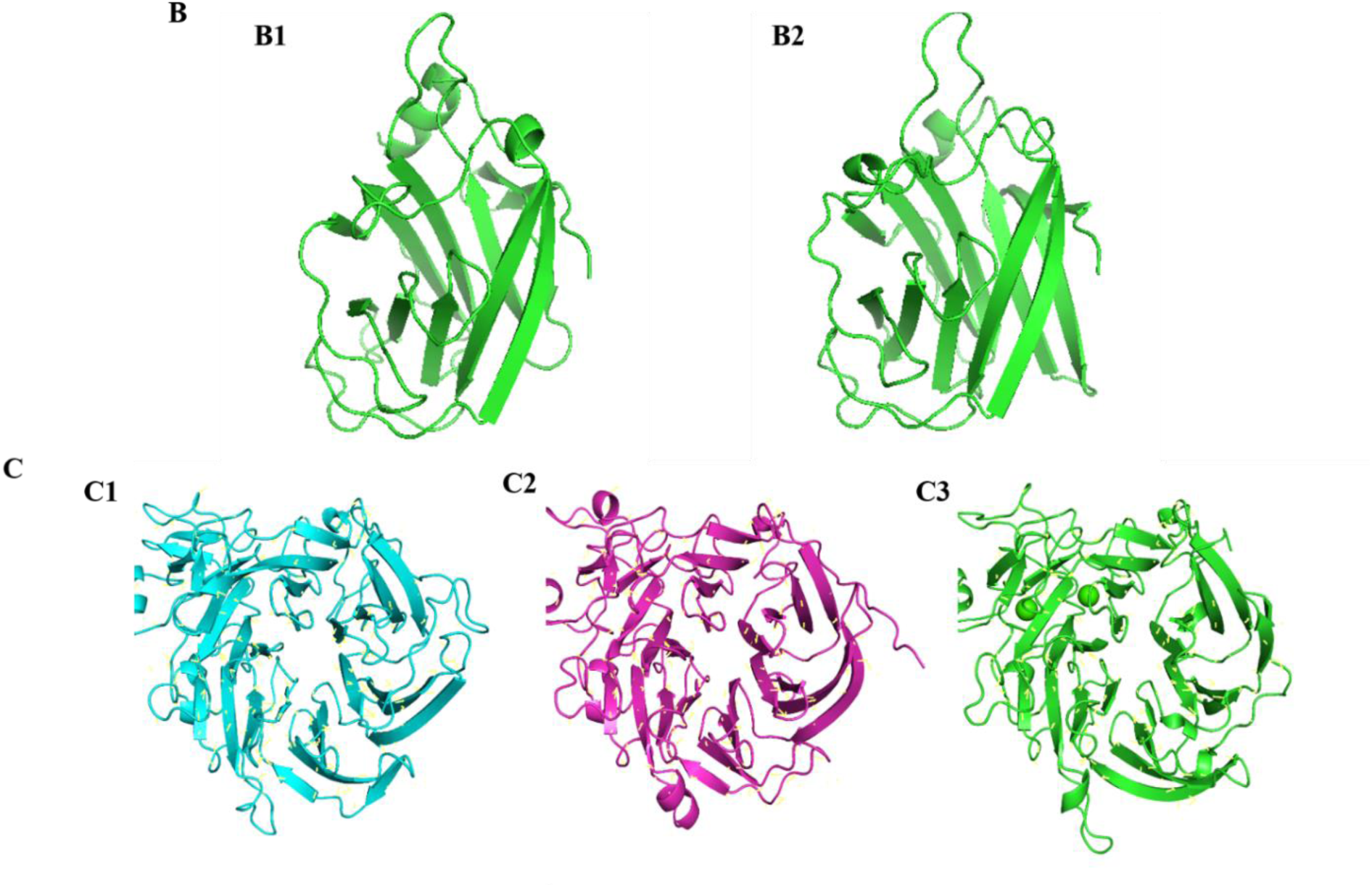
(A) Phylogenetic tree analysis of AA12 PQQ-dependent oxidoreductases. Red: *Tth*AA12A, Green: *Tth*AA12B, Blue: *Tte*AA12A, Purple: *Tr*AA12, Fuchsia: *Cc*PDH. (B) Structural models of *Tth*AA8A (B1) and *Tth*AA8B (B2). (C) Structural models of *Tth*AA12A (C1), *Tth*AA12B (C2), and *Tte*AA12A (C3) obtained from SWISS-MODEL.

Furthermore, the enzymes studied in this work display a distant evolutionary relationship from previously characterized AA12 enzymes. The amino acid sequences of AA12 enzymes and AA8 modules were submitted to the AlphaFold 2 and SWISS-MODEL web servers for 3D structural model simulation. The quality of the resultant models was assessed using QMEANDisCo. Ultimately, the SWISS-MODEL outputs were selected due to their higher scores (0.71 ± 0.05) compared to AlphaFold 2 (0.68 ± 0.05). AA12 enzymes and AA8 modules was compared with crystallized enzymes by Dali server (http://ekhidna2.biocenter.helsinki.fi/dali/). The results revealed that the three AA12 enzymes including *Tth*AA12A, *Tth*AA12B and *Tte*AA12A exhibited the highest structural similarity to the AA12 domain of CcPDH, while the AA8 modules demonstrated the closest structural resemblance to the AA8 domain of *Nc*CDH (PDB ID: 4QI7) from *Neurospora crassa* OR74A (Figure 1B and 1C). Specifically, when comparing the SWISS-MODEL-derived 3D structure of AA12 enzymes with *Cc*PDH (PDB ID: 6JWF), we found that their core structures were composed of six β-sheets arranged in a helical formation (3, 14). Multiple sequence alignment revealed that the active sites, PQQ binding sites, and calcium ion binding sites of AA12 enzymes were nearly identical (Figure S2 and Table S1). These sites were highly conserved within the AA12 family, suggesting that catalysis and substrate binding mechanisms may be similar across these enzymes (3, 14). Multiple sequence alignment revealed that *Tth*AA8A and *Tth*AA8B showed 44.34% identity with the AA8 domains in *Cc*PDH, *Mt*CDH from *Myriococcum thermophilum*, *Cc*CDH from *N. crassa* and *Pc*CDH from *Phanerochaete chrysosporium* CDH (Figure S3), and the heme-binding residues (Met and His) are highly conserved (26, 7, 27).

### Expression and purification of AA12 enzymes and AA8 modules in *P. pastoris* X-33

Three AA12 enzymes and two AA8 modules were successfully expressed in *P. pastoris* X-33 and purified via Ni-NTA chromatography. The theoretical molecular weights of *Tth*AA12A, *Tte*AA12A and *Tth*AA12B were predicted as 45.7 kDa, 43.9 kDa and 48.8 kDa, respectively, using ExPASy ProtParam (https://web.expasy.org/protparam/, accessed on 1 April 2025). SDS–PAGE analysis revealed that *Tth*AA12A and *Tte*AA12A displayed a noticeable tail with the experimental molecular weight of approximately 50 kDa, suggesting potential glycosylation modification (Figure 2). While *Tth*AA12B displayed single band with the experimental molecular masses of approximately 44 kDa, close to its theoretical molecular weight. Both *Tth*AA8A and *Tth*AA8B have the molecular weight of approximately 25 kDa, close to their predicted theoretical sizes (Figure 2). The reconstitution of the apoenzymes of purified *Tth*AA12A, *Tth*AA12B and *Tte*AA12A was assayed by measuring the enzyme activity of PQQ-saturated AA12 enzymes. The maximal activity was observed for the AA12 enzymes with 60-minutes PQQ saturation (Figure S4). The PQQ occupancy in AA12 enzymes was also determined using the TCA method. Quantitative analysis revealed that the PQQ concentration closely matched the protein concentration, suggesting complete saturation of the cofactor binding sites (Table S2). Furthermore, the supplementation with 1 mM calcium chloride did not significantly alter AA12 enzyme activity compared to that of no addition, implying that the calcium ion was also fully saturated under the experimental conditions. Thus, the AA12 enzymes with a saturation time of 60 minutes was used in the subsequent experiments (Table S3).

**Figure 2.**
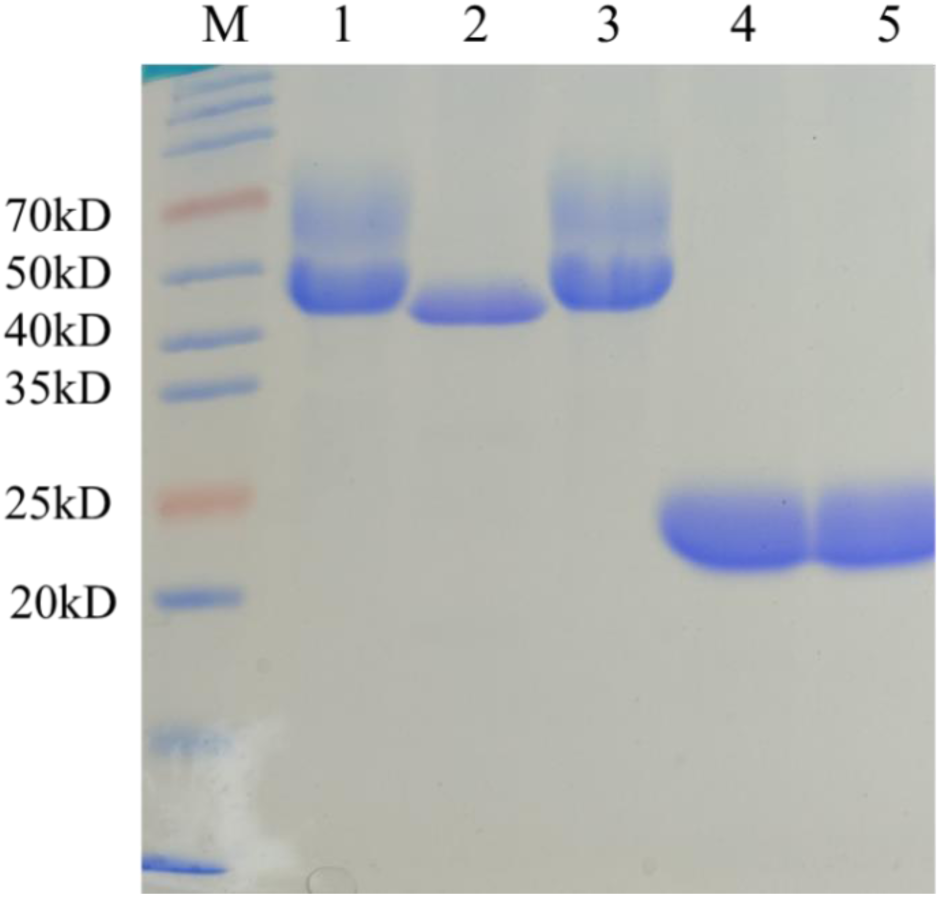
SDS-PAGE analysis of the purity of AA12 enzymes and AA8 modules: M: molecular weight marker; lane 1, *Tth*AA12A; lane 2, *Tth*AA12B; lane 3, *Tte*AA12A; lane 4, *Tth*AA8A; lane 5, *Tth*AA8B.

### Effects of pH, temperature on AA12 enzymes activity

Effects of pH and temperature on AA12 enzymes activity were measured using L-fucose as substrate and DCIP as electron acceptor across different pHs and temperatures. The optimal temperatures for *Tth*AA12A, *Tth*AA12B, *Tte*AA12A were 60°C, 55°C and 60°C, respectively (Figure 3A). Notably, *Tth*AA12A and *Tte*AA12A demonstrated superior thermophily, retaining >50% relative activity at 70°C compared to *Tth*AA12B. After incubated at 30°C for 72 h, approximately 50% residual activities were remained for *Tth*AA12A and *Tte*AA12A, but only 30% for *Tth*AA12B.

**Figure 3.**
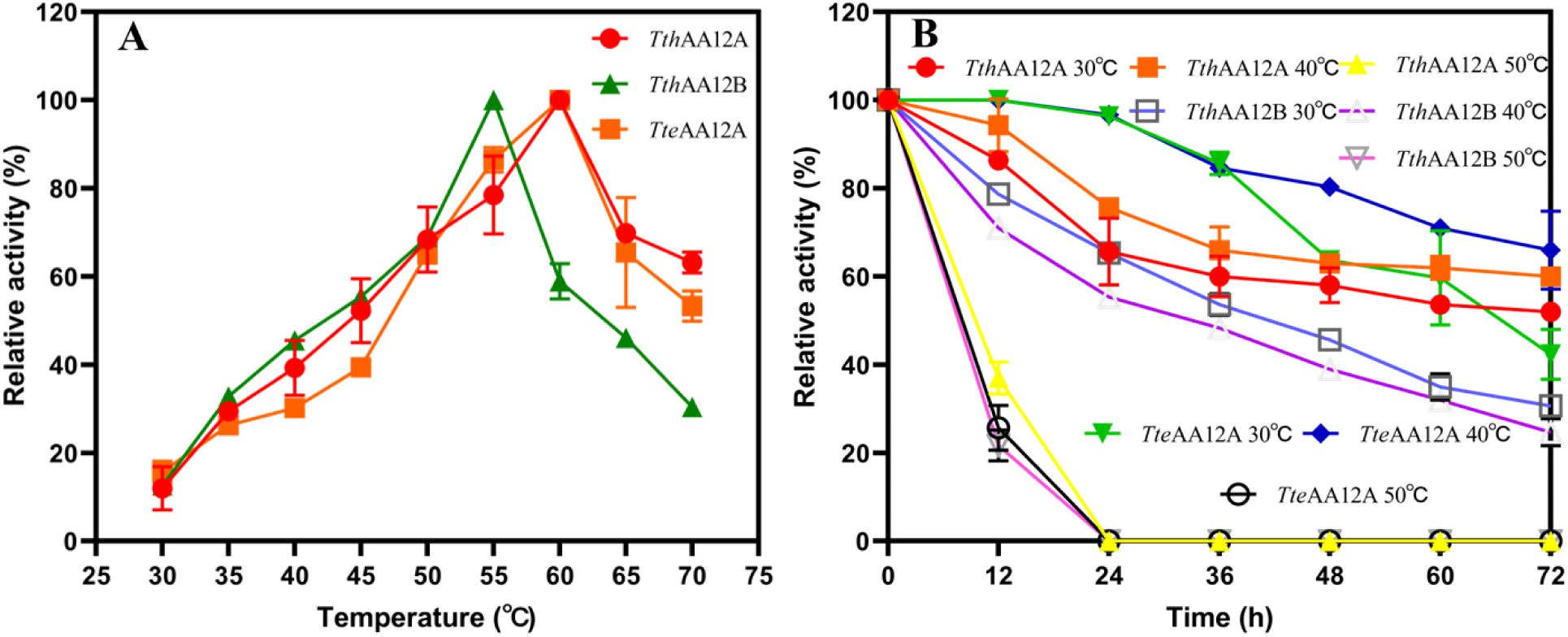
Effects of temperature on the reduction of DCIP by AA12 enzymes. (A) Optimal temperatures of AA12 enzymes using L-Fucose as the substrate. (B) Thermostability of AA12 enzymes using L-Fucose as the substrate and DCIP as the electron acceptors.

Stability at 40°C was comparable to 30°C, while 50°C exposure caused rapid inactivation within 24 h, though >25% activity persisted during the first 12 h (Figure 3B). All AA12 enzymes preferred neutral to weakly acidic conditions, with optimal pH values of 6.0 (*Tth*AA12A), 7.0 (*Tth*AA12B), and 6.5 (*Tte*AA12A), respectively (Figure 4A). They demonstrated broad pH stability, maintaining >20% activity after 72 h at pH 4.0-10.0 and >50% activity at pH 6.0-9.0 (Figure 4B, 4C and 4D).

**Figure 4.**
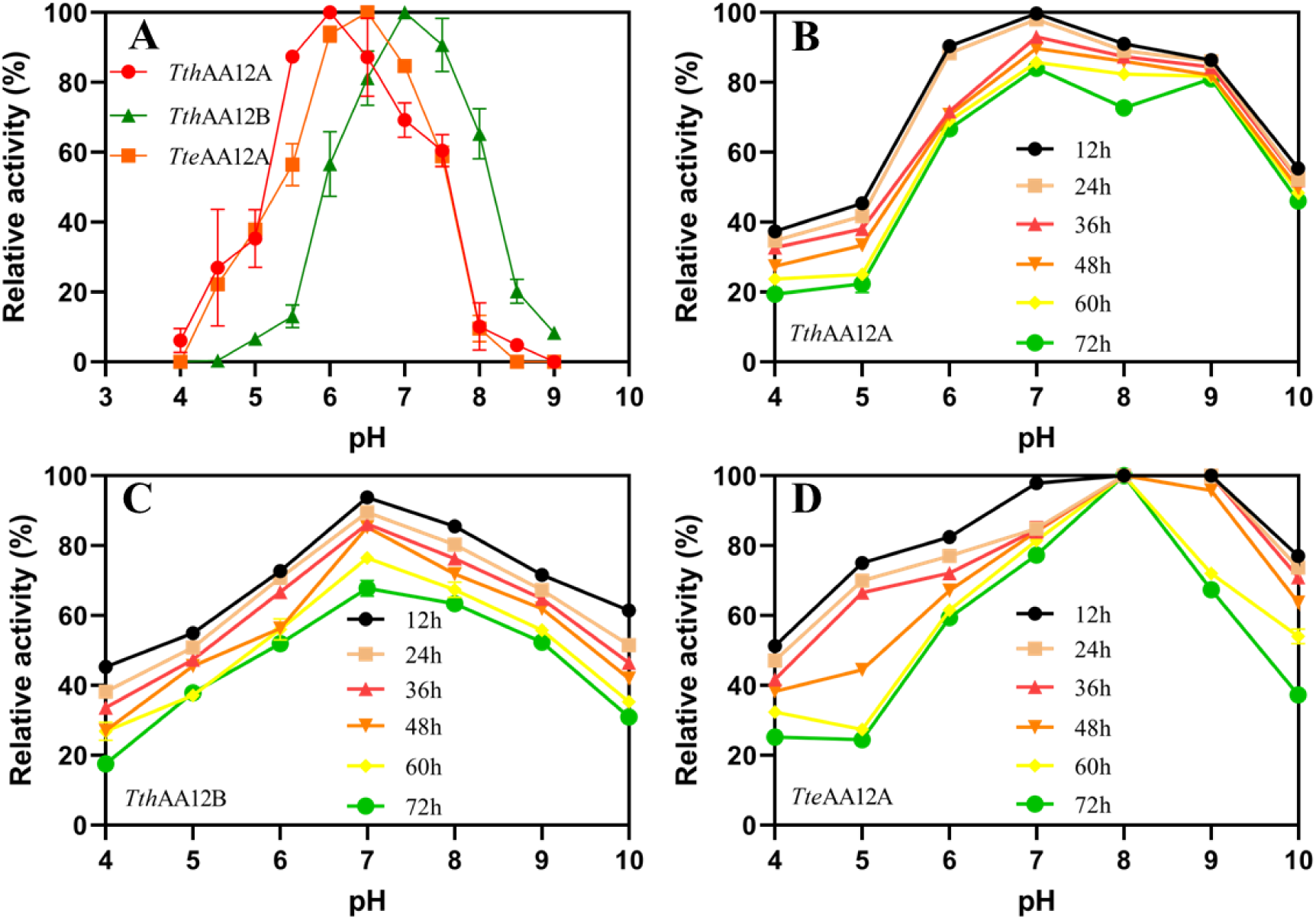
Effects of pH on the reduction of DCIP by AA12 enzymes using L-fucose as the substrate. (A) Optimal pHs of AA12 enzymes. (B, C, D) pH stability of *Tth*AA12A, *Tth*AA12B and *Tte*AA12A, respectively.

### Substrate specificity and kinetics of AA12 enzymes

Among all tested substrates, the three AA12 enzymes exhibited catalytic activity exclusively toward three rare sugars including L-fucose, L-ribose, and D-lyxose. As shown in Table 1 and Figure S5, all three AA12 enzymes demonstrated high *K*_m_ values (>100 mM) for their active substrates, indicating weak substrate affinity. Specifically, the previous characterized *Cc*PDH and *Tr*AA12 showed a *K*_m_ of 24.8 mM and 99 mM for L-fucose, respectively, while the corresponding values for *Tth*AA12A, *Tth*AA12B and *Tte*AA12A are 287 mM, 423 mM and 459 mM, respectively. *Tth*AA12A, *Tth*AA12B and *Tte*AA12A exhibited the highest catalytic efficiency for D-lyxose, L-ribose and L-fucose in terms of *K*_cat_/*K*_m_ (0.08 s⁻¹·mM⁻¹, 0.026 s⁻¹·mM⁻¹ and 0.03 s⁻¹·mM⁻¹, respectively), collectively indicating lower catalytic efficiency than *Cc*PDH but much higher than *Tr*AA12. Molecular docking of three AA12 enzymes with L-fucose using Schrödinger software revealed hydrogen bond interactions between them. Compared to *Cc*PDH, AA12 enzymes showed relatively less hydrogen bonds and a higher free energy (Figure 5).

**Figure 5.**
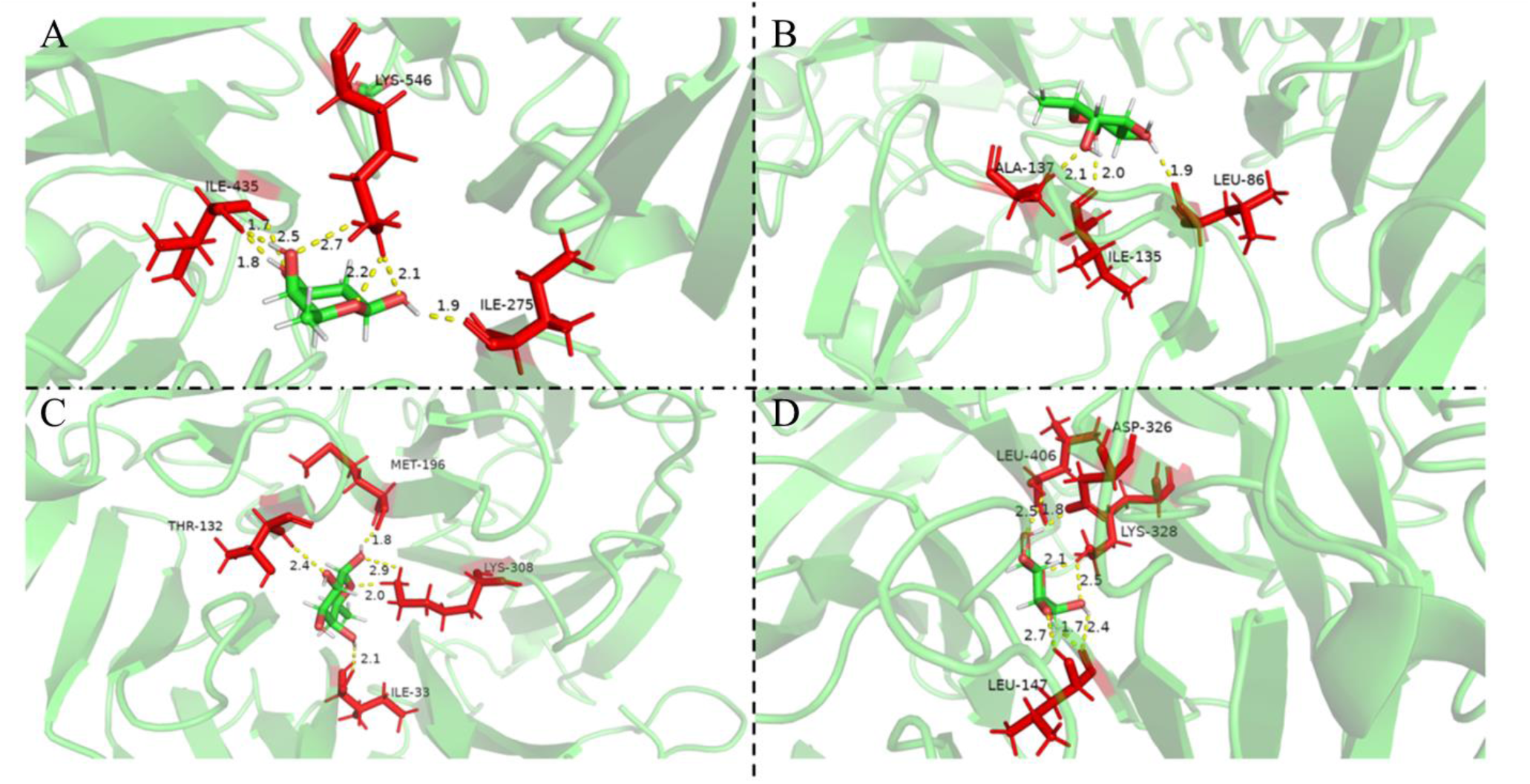
Visualization results of molecular docking between four AA12 enzymes and L-Fucose in PyMOL. (A, B, C, D) For *Cc*PDH, *Tth*AA12A, *Tth*AA12B, and *Tte*AA12A, respectively. Green: PQQ; Red: amino acid residues interacting with PQQ; Yellow dashed lines: hydrogen bonds.

**Table 1.** Kinetic constants of the tested AA12 enzymes and previously characterized AA12.

| | | $K_m$ (mM) | $K_{cat}$ ($\text{s}^{-1}$) | $K_{cat}/K_m$ ($\text{s}^{-1}\text{ mM}^{-1}$) | Reference |
| --- | --- | --- | --- | --- | --- |
| L-Fucose | ***TthAA12A*** | 287 | 12.66 | 0.04 | This study |
|  | ***TthAA12B*** | 423 | 5.21 | 0.01 | This study |
|  | ***TteAA12A*** | 459 | 12.73 | 0.03 | This study |
|  | ***CcPDH*** | 24.8 | 56.4 | 2.27 | [5] |
|  | ***TrAA12*** | 99 | 0.012 | 0.119 | [18] |
| D-Lxyose | ***TthAA12A*** | 158 | 12.77 | 0.08 | This study |
|  | ***TthAA12B*** | 908 | 5.03 | 0.005 | This study |
|  | *TteAA12A* | 495 | 11.07 | 0.022 | This study |
|  | *CcPDH* | 66.8 | 12.9 | 0.19 | [5] |
| L-Ribose | *TthAA12A* | 449 | 11.08 | 0.024 | This study |
|  | *TthAA12B* | 265 | 6.79 | 0.026 | This study |
|  | *TteAA12A* | 469 | 3.86 | 0.008 | This study |

### Transient-State kinetic studies

The successive PQQ reduction in AA12 enzymes was monitored at 330 nm by stopped-flow spectroscopy during substrate oxidation. As shown in Figure 6A, 6B and 6C, all AA12 enzymes displayed comparable PQQ reduction rate constants (*K*_obs330_) at equivalent substrate concentrations. The values of *K*_obs330_ (s^-1^) are 1.048, 0.875 and 0.758 for *Tte*AA12A, *Tth*AA12A and *Tth*AA12B, respectively. To examine potential IPET between AA12 enzymes and AA8 modules, we measured heme b reduction rate in AA8 modules at 563 nm at the presence of AA12 enzymes, a process dependent on PQQ-to-heme electron transfer. While characteristic exponential curves were absent, *Tth*AA8B showed significant absorption increases by AA12 enzymes (Figure 6D), suggesting weak electron transfer between them. In contrast, no absorption increase was observed for *Tth*AA8A. Although the precise mechanism remains unclear, these observations imply that the electron transfer occurs between AA12 enzymes and *Tth*AA8B. In comparison, the inter-protein electron transfer rate between single-domain AA12 enzymes with free AA8 modules is obviously lower than inter-domain between AA3 DH domains and AA8 domains in CDHs (28–30).

**Figure 6.**
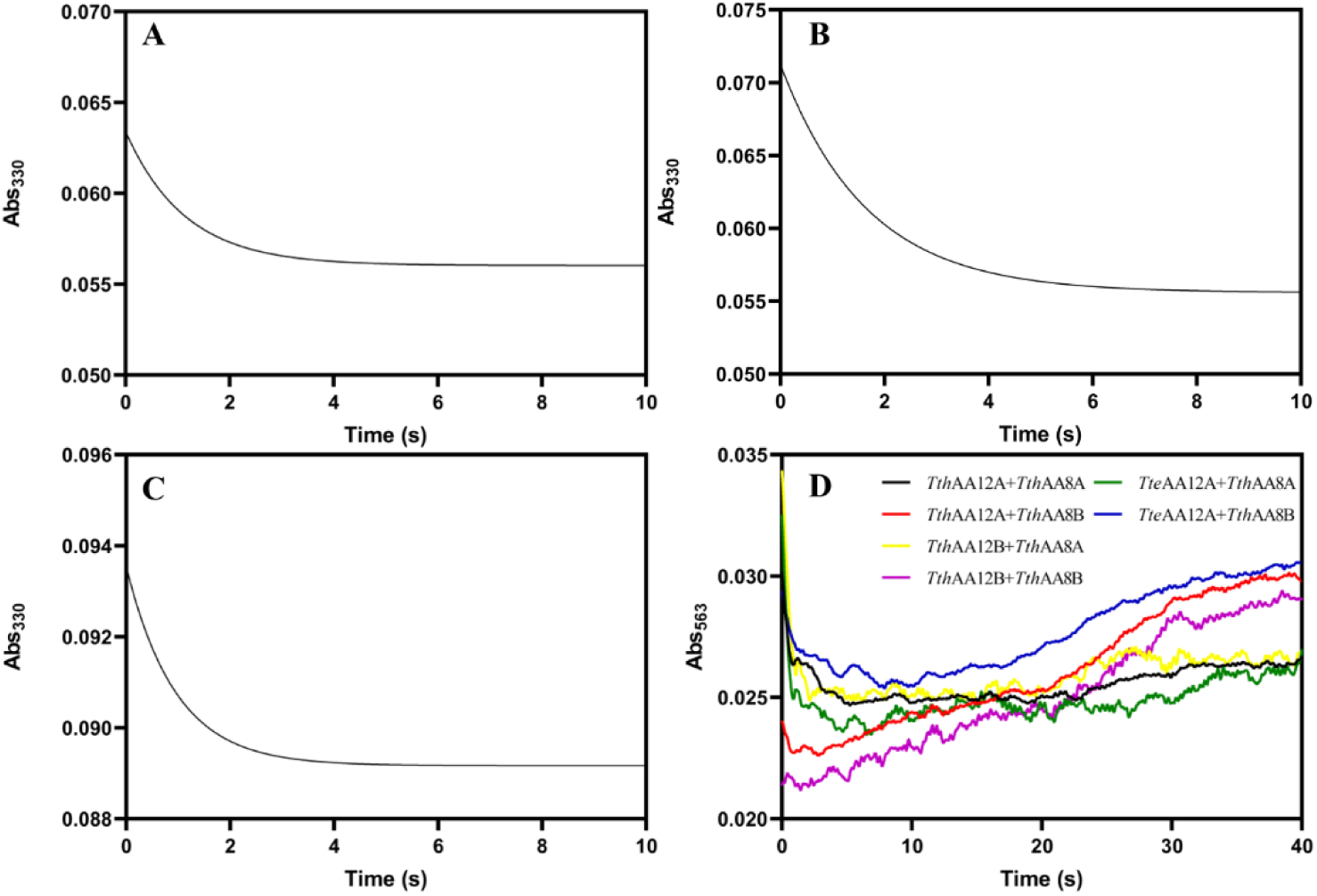
Reduction curves of PQQ in AA12 enzymes of *Tth*AA12A (A), *Tth*AA12B (B) and *Tte*AA12A (C), and heme b reduction curves in AA8 modules (k_obs563_) by AA12 enzymes (D). Black: *Tth*AA12A + *Tth*AA8A, red: *Tth*AA12A + *Tth*AA8B, yellow: *Tth*AA12B + *Tth*AA8A, purple: *Tth*AA12B + *Tth*AA8B, green: *Tte*AA12A + *Tth*AA8A, blue: *Tte*AA12A + *Tth*AA8B.

### UV-Vis spectroscopy confirms typical AA8 features and electrons transfer between AA12 enzymes and AA8 module

The above stopped-flow experiments suggested a certain degree of existence of the electron transfer between AA12 enzymes and AA8 modules *Tth*AA8B, but it may not occur between AA12 enzymes and AA8 modules *Tth*AA8A. So, we used a UV-Vis spectrophotometer to investigate the changes of the characteristic peaks of the AA8 modules before and after reduction action by dithionite (Figure 7A). The spectral properties of AA8 modules were studied by recording UV-Vis spectra ranging from 400 nm to 600 nm. A typical cytochrome spectrum was observed for *Tth*AA8B, with the dominant heme b Soret band with a maximum at 420 nm. Upon the addition of dithionite this band shifted to 429 nm due to the reduction of heme b in *Tth*AA8B, while the reduction characteristic peaks at 533 nm and 563 nm was significantly increased (Figure 7A1). However, *Tth*AA8A did not display this characteristic peak before and after dithionite reduction, indicating that it does not have heme b or has extremely low proportion (Figure 7A2) of the occupied cofactor in binding site. This observation may explain the no obvious absorption increase at 563 nm of *Tth*AA8A in stopped-flow experiment (Figure 6D). Furthermore, the time-dependent electron transfer between AA12 enzymes and *Tth*AA8B was investigated by UV-Vis spectrophotometer. The height of the characteristic peaks of heme b at 533 nm and 563 nm increased as reaction time increasing, indicating that the electrons can be transferred from reduced AA12 enzymes after substrate oxidation to *Tth*AA8B (Figure 7B). Due to the highly efficient interaction between *Tth*AA12A and *Tth*AA8B, *Tth*AA8B was fully reduced within 15 minutes, equivalent in efficiency to dithionite (Figure 7B1), while *Tth*AA8B could be fully reduced by *Tth*AA12B and *Tte*AA12B under same condition (Figure 7B2 and 7B3).

**Figure 7.**
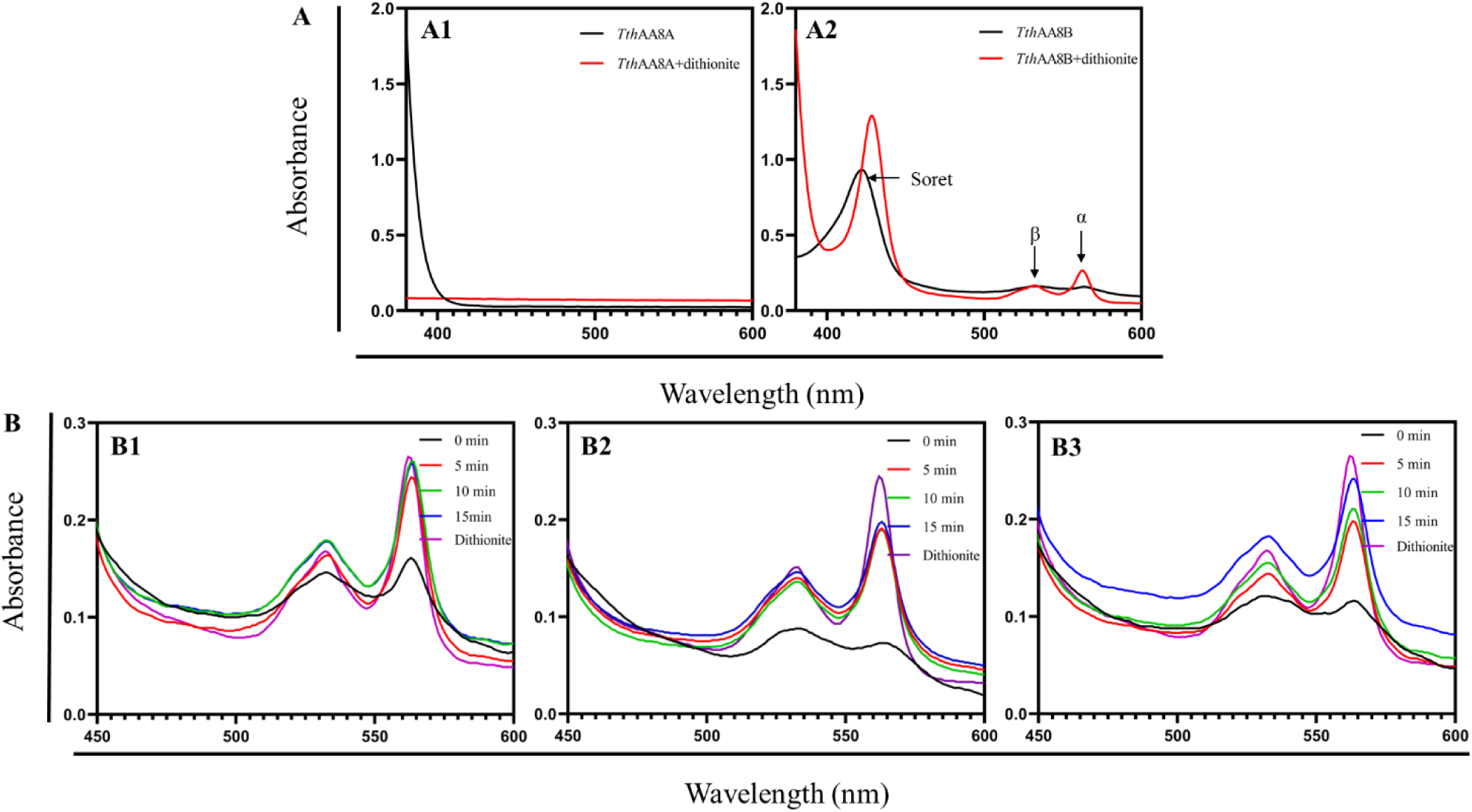
(A) Spectral analysis of *Tth*AA8A (A1) and *Tth*AA8B (A2) in oxidized (black) and dithionite-reduced (red) states. The absorption maxima of the Soret-, β-and α-heme band are indicated. (B) Spectral analysis of *Tth*AA8B reduced by *Tte*AA12A (B1), *Tth*AA12B (B2) and *Tte*AA12A (B3). *Tth*AA8B and AA12 enzymes were mixed in equimolar amounts of 10 µmol L^−1^ in MOPS buffer (pH 6.0). The reaction was started after addition of 100 mmol L^−1^ L-Fucose at 45°C and spectra recorded with 5-mins intervals. Black line: oxidized *Tth*AA8B; Red line: after 5 mins reaction; Green line: after 10 mins reaction; Blue line: after 15 mins reaction; Purple line: fully reduced by sodium dithionite.

### Oxidase activity and hydrogen peroxide production

Monitoring H_2_O_2_ production rates at 1 μM AA12 enzymes over 20 minutes revealed that all three AA12 enzymes generated substantial H_2_O_2_ through oxidase activity, with *Tth*AA8B exhibiting the fastest oxidation rate, followed by *Tth*AA12A and *Tte*AA12A (Figure 8A). The oxidase activities of AA12 enzymes, measured using the HRP/Amplex Red assay, were 0.013±0.003, 0.017±0.002 and 0.013±0.002 U/mg for *Tth*AA12A, *Tth*AA12B and *Tte*AA12A, respectively. Their dehydrogenase activities were 16.498±1.188, 17.335±2.003 and 21.396±3.403 U/mg, respectively. In comparison, AA12 enzymes exhibited high dehydrogenase activity and weak oxidase activity. Of note, the addition of *Tth*AA8B reduced H_2_O_2_ generation rates by approximately three-fold likely as external electron receptor for AA12 enzymes (Figure 8B).

**Figure 8.**
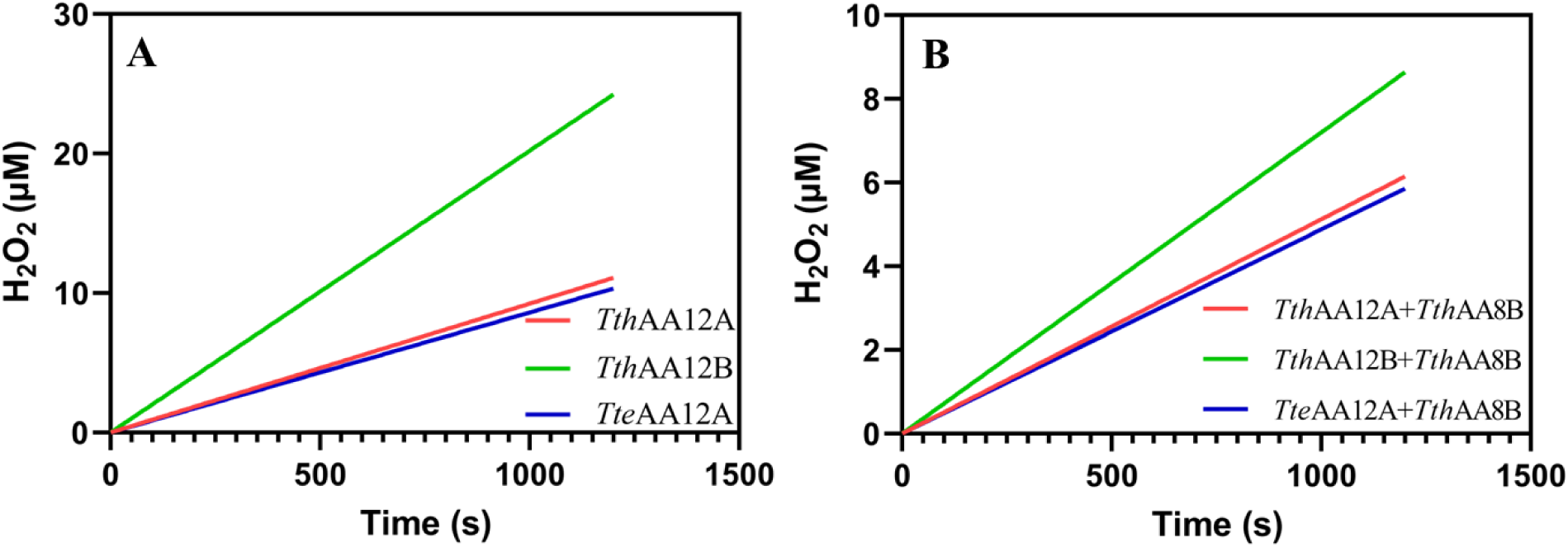
The H₂O₂ production curves of AA12 enzymes. (A) Without the addition of *Tth*AA8B. (B) With the addition of *Tth*AA8B.

### Activation of *Nc*LPMO9C by AA12 enzymes alone or cooperatively with AA8 modules

In the AA12 enzyme-driven *Nc*LPMO9C reactions, excess L-fucose (200 mM) was added as the substrate for AA12 enzymes, and the activities of AA9 LPMOs on cellulose driven by either AA12 enzyme alone or combined AA12 enzymes and AA8 module were quantified in terms of the sum of C4-oxidized product peak areas. As shown in Figure S6 and Figure 9, three AA12 enzymes alone were all capable of activating *Nc*LPMO9C to generate oxidized products from PASC, with the amount of the oxidized products increased as increase of reaction time. No oxidized products were observed in either the heat-inactivated enzyme or the PQQ control group (Figure S6). When using sodium dithionite-reduced *Tth*AA8B as the sole electron donor (with equimolar concentrations of AscA as control), distinct *Nc*LPMO9C oxidation product peaks were observed in the 100 μM reduced *Tth*AA8B-implemented samples (Figure S7). The oxidized product peak area in 200 μM reduced *Tth*AA8B-implemented samples approximately doubled compared to that of 100 μM. This result demonstrated that the sodium dithionite-reduced *Tth*AA8B can also transfer electrons to *Nc*LPMO9C to activate *Nc*LPMO9C reaction. The activation of *Nc*LPMO9C by the combined AA12 enzymes and AA8 module *Tth*AA8B was also investigated in parallel (Figure S6). As expected, the oxidized products with addition of *Tth*AA8B were significantly increased than those without the addition of *Tth*AA8B after 12h and 24h reactions, implying that *Tth*AA8B could facilitate the driving effect of AA12 enzymes on *Nc*LPMO9C activity (Figure 9). Correspondingly, the addition of *Tth*AA8B in *Tth*AA12A-, *Tth*AA12B- or *Tte*AA12A-driven *Nc*LPMO9C system generated 74% and 71.6%, 116% and 25.7%, or 101% and 24% higher oxidized products than that of without addition of *Tth*AA8B after 12h and 24h reaction, respectively (Figure 9).

**Figure 9.**
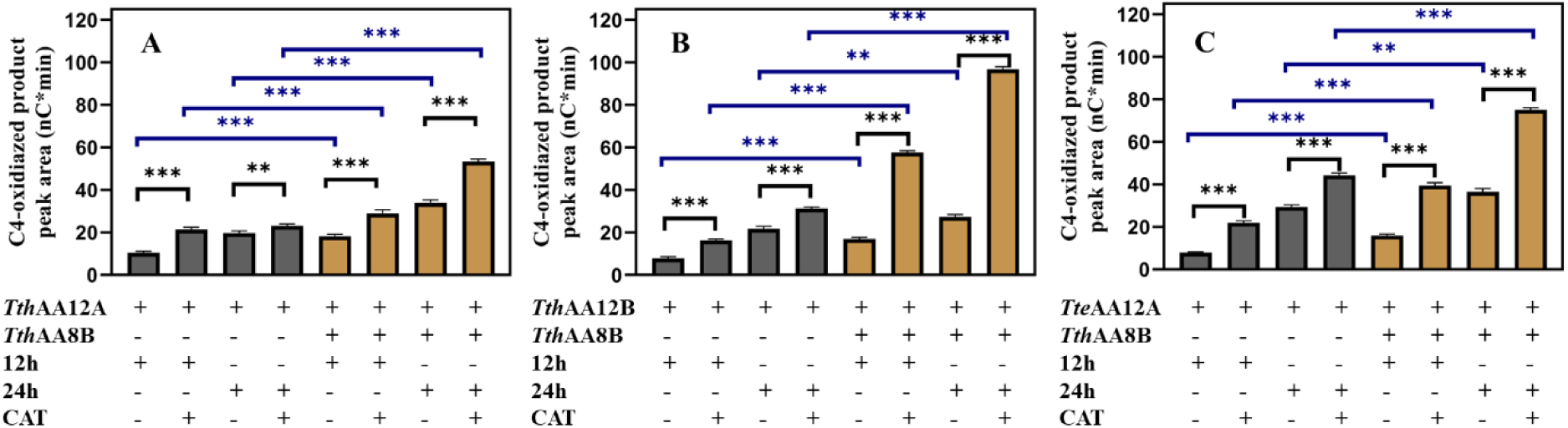
Comparative analysis of oxidized product levels of *Nc*LPMO9C driven by three AA12 enzymes alone or cooperatively with AA8 module. (A) *Tth*AA12A-driven *Nc*LPMO9C oxidative degradation of PASC. (B) *Tth*AA12B-driven *Nc*LPMO9C oxidative degradation of PASC. (C) *Tte*AA12A-driven *Nc*LPMO9C oxidative degradation of PASC. **Gray**: AA12-driven reaction without *Tth*AA8B; **Brown**: AA12-driven reaction with *Tth*AA8B

Since AA12 enzymes display oxidase activity and could generate hydrogen peroxide, we further analyzed the driving efficiency of AA12 enzymes on *Nc*LPMO9C activity in the presence of CAT (Figure S8). When added 10 µg CAT to the system, the amounts of oxidative products generated either by AA12 enzymes-*Nc*LPMO9C system or AA12 enzymes/AA8 module-*Nc*LPMO9C system were significantly increased by approximately two- to three-fold compared to these without addition of CAT (Figure 9). From above studies, it is convincible that three novel single-domain AA12 PQQ-dependent oxidoreductases (*Tth*AA12A, *Tth*AA12B and *Tte*AA12A) are capable of directly and cooperatively driving LPMO activity with AA8 modules. This driving effect may not be due to the generation of hydrogen peroxide by the oxidase activity of AA12 enzymes.

### EPR spectroscopy

EPR spectroscopy was used to investigate whether AA12 enzymes can directly transfer electrons to LPMO and reduce the divalent copper of LPMO to monovalent copper ions. The extent of LPMO-Cu(II) reduction to Cu(I) was monitored as a decrease of the Cu(II) EPR signal due to the silence of LPMO-Cu(I) species in EPR (31). The addition of AA12 enzymes to *Nc*LPMO9C pre-incubated with L-fucose under anaerobic conditions led to a decrease (40–80%) in *Nc*LPMO9C-Cu(II) signal (Figure 10). Despite only partial reduction of the Cu(II) center was observed likely due to kinetic or thermodynamic barriers, our data demonstrated that three one-module AA12 enzymes confers direct electron transfer from the AA12-bound PQQ to the active site of the LPMO and reduce the divalent copper of LPMO to monovalent copper ions.

**Figure 10.**
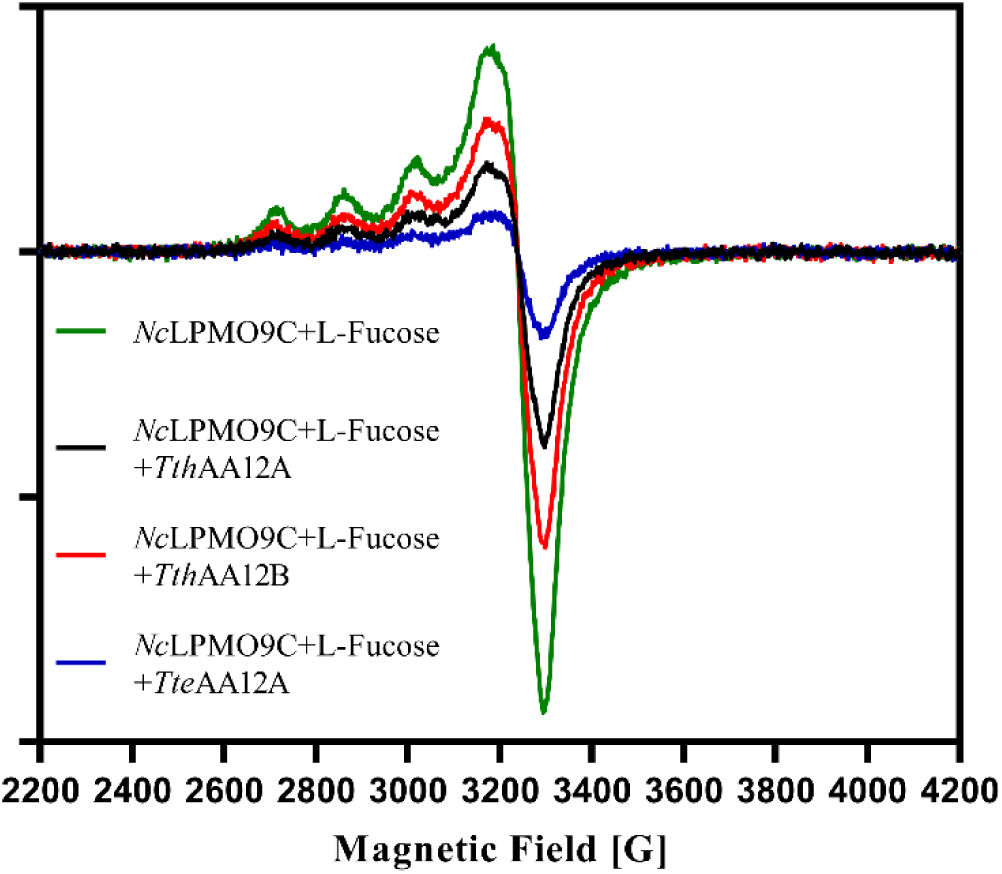
EPR spectra of NcLPMO9C−Cu(II) (20 μM) in buffer (blue line), in the presence of AA12 enzymes. All EPR solutions and experiments were performed under anaerobic conditions. Samples were in 50 mM MOPS buffer, pH 6.5 and spectra were recorded at 77 K with a 19.73 mW microwave power and a 3 mT modulation amplitude.

## DISCUSSION

The fungal AA12 PQQ-dependent oxidoreductases are widely distributed in cellulolytic fungi. Previous studies have confirmed that the multi-domain oxidoreductase *Cc*PDH could act as a reductant for LPMOs for the redox-mediated oxidative cleavage of cellulose (11). The co-occurrence of the AA12 and the AA8 domains in *Cc*PDH is essential for electron transfer between AA12 domain and AA8 domain, and AA12 domain and LPMO (11). However, a search on the CAZy database reveals that only a few members of the AA12 family sequences exist in a multi-domain form connected to the AA8 module via a flexible linker peptide, in contrast, the majority of AA12 sequences in fungi are nonmodular. Meanwhile, a considerable proportion of members in the AA8 family are single-domain as well. With aims to investigate the auxiliary functional roles of single-domain AA12 enzymes in cellulolytic fungi, three of four genes encoding for AA12 oxidoreductase from well-known cellulolytic fungi *T. thermophilus* ATCC 42464 and *Thermothielavioides terrestris* NRRL 8126 were successfully expressed and characterized by their main biochemical and physiochemical properties in this study. The interplay between single-domain AA12 enzymes and AA8 module, as well as single-domain AA12 enzymes and *Nc*LPMO9C, were comprehensively monitored by visible absorption spectral scanning, stopped-flow experiment, and EPR. Additionally, the auxiliary roles of AA12 enzymes or/and AA8 modules in driving LPMO reaction was also demonstrated by the C4-oxidizing *Nc*LPMO9C using PASC as cellulosic substrate.

Our works revealed that three tested single-domain AA12 enzymes typically exhibit dehydrogenase activity towards L-fucose, L-ribose, and D-lyxose, similar as *Cc*PDH (1, 2). In comparison, three single-domain AA12 enzymes displayed narrower substrate specificities with lower substrate affinity and catalytic efficiency than *Cc*PDH, despite both of them display highly similar predicted structure and co-factor PQQ was fully occupied (1,2). Hydrogen bonds play a crucial role in stabilizing protein secondary structures and maintaining conformational stability (32, 33). The more hydrogen bonds between *Cc*PDH and L-fucose were found in the subpocket by molecular docking, which may improve protein stability and substrate binding affinity, potentially contributing to increased catalytic activity of *Cc*PDH (9,32,33). Surprisingly, though three tested single-domain AA12 enzymes displayed dehydrogenase activity higher than previously characterized single-domain *Tr*AA12 from *T. reesei* (14), all three tested single-domain AA12 enzymes were able to use oxygen as electron acceptor (Figure 8), however, they reduced molecular oxygen much less efficiently than DCIP (with an approximately ratio of 1000:1 of dehydrogenase/oxidase activity). So, these tested single-domain AA12 enzymes should be considered dehydrogenases with trace residual oxidase activity. So far, only two PQQ-dependent sugar dehydrogenases were characterized, but their oxidase activity was not reported (1, 2, 14). The residual oxidase activity was detected in some FAD-dependent dehydrogenases such as AA3-1 CDHs (34–36) and AA3-2 aryl-alcohol:quinone oxidoreductases (*Pc*AAQO2 and *Pc*AAQO3) (31,37). The distinctively high dehydrogenase/oxidase activity of all three tested single-domain AA12 enzymes in this study raises the question whether other AA12 PQQ-dependent oxidoreductases also has week oxidase activity. More fungal AA12 oxidoreductases need to be characterized to explore whether and to what extent other fungal AA12 enzymes have residual oxidase activity.

The activation of fungal LPMOs requires an electron donor to reduce Cu^2+^ to Cu^+^ and utilizes O_2_ or H_2_O_2_ as co-substrates to drive catalytic activity (7, 8). The electron donors can be small molecule reductants, such as ascorbic acid, or flavoproteins like CDHs (7, 8). Compared to using small molecule reductants as electron donors, it was considered that employing an enzyme redox partner as electron donor is advantageous because this enables a kinetically controlled supply of electrons to the LPMO (7, 8). The established enzymatic redox partners, including CDHs and *Cc*PDH, exhibit multi-domain structural architectures, and their electron transfer function relays on the appended AA8 cytochrome b domain (7,11). Significant progresses have been made in developing and engineering multi-domain redox enzymes for industrial applications (9, 38). However, research investigating the role of single-domain redox enzymes and free cytochrome modules for lytic polysaccharide monooxygenases (LPMOs) remains limited. So far, few single-domain flavoenzymes were discovered to drive lytic polysaccharide monooxygenases for oxidative degradation of cellulose. These proteins include AA3_2 aryl-alcohol quinone oxidoreductases from *Pycnoporus cinnabarinus* (31), a novel AA7 cellooligosaccharide dehydrogenase *Fg*CelDH7C from *F. graminearum* (15). All three single-domain flavoenzymes exhibit typical characteristics of dehydrogenase, despite AAQO2 displays residual oxidative activity (15, 31). However, not all of FAD-containing dehydrogenase are capable of driving LPMO reaction. For example, *Mo*Chi7A that possesses a low oxidase activity on cellooligosaccharides did not fuel LPMO activity (15). AAQO2 with residual oxidase activity also exhibited lower driving efficiency than the strict dehydrogenase AAQO1 (31). It was reported that an aryl-alcohol oxidase (*Tt*AAOx) from *T. thermophilus* in the presence of veratryl alcohol could drive the LPMO activity, however, no direct electron transfer between the enzymes was detected, indicating that *Tt*AAOx predominantly worked with *Tt*LPMO9H as a provider of H_2_O_2_ (39). The current study demonstrated that all three tested single-domain AA12 enzymes, even without a cytochrome domain, could directly transfer the electrons to Cu(II) in *Nc*LPMO9C after oxidizing the substrate, thereby reducing Cu(II) into Cu(I) and activating LPMOs for efficient cellulose degradation. We excluded the interference from free PQQ. This observation led us to hypothesize that all members of the single-domain AA12 family PQQ-dependent dehydrogenases may transfer electrons to LPMOs and activates LPMOs for efficient cellulose degradation, since they are prevalent in cellulolytic fungi. More single-domain AA12 enzymes need to be characterized to answer this hypothesis in future work.

The deletion of the cytochrome domain in multi-domain *Cc*PDH eliminated the fueling of LPMO activity (11). Similarly, the deletion of the cytochrome domain in multi-domain CDH impaired the priming of the LPMO (36), indicating that the active site topology of the dehydrogenase domain in multi-domain AA12 or AA3 redox enzymes may be evolved to satisfy the interaction with the appended AA8 cytochrome b domain. In contrast, the electron transfer and the interaction of AA7 cellooligosaccharide dehydrogenase with the LPMO is likely to occur at the substrate-binding face (si-side) of the FAD, which is consistent with the rather open active site topology of *Fg*CelDH7C (15). Given that single-domain AA12 enzymes alone can act as electron donor for LPMOs and the prevalence of free AA8 modules, we questioned the evolutionary significance of free AA8 module and its interaction with single-domain AA12 enzyme and LPMOs in biodegradation of lignocellulose. We hypothesized that, similar to its role in multi-domain enzymes, the free AA8 modules might serve as an electron acceptor for the single-domain AA12 enzymes and transfer electrons to terminal acceptors. This hypothesis was supported in this study through two key findings. First, UV-Vis spectral scanning and stopped-flow experiments confirmed the *Tth*AA8B’ electron acceptance from AA12 enzymes-mediated substrate oxidation (Figure 6D and Figure 7B), Second, dithionite-reduced *Tth*AA8B effectively activated *Nc*LPMO9C, demonstrating its capacity as electron donor (Figure S8). Furthermore, the presence of *Tth*AA8B effectively reduced hydrogen peroxide compared to AA12 enzymes alone (Figure 8B). As expected, the addition of AA8 module *Tth*AA8B into the AA12 enzymes-*Nc*LPMO9C system significantly enhanced *Nc*LPMO9C oxidative product yields (Figure 9), which may be attributed to the acceleration of electron transfer from AA12 enzymes to *Nc*LPMO9C and the attenuation of H_2_O_2_ accumulation mediated by *Tth*AA8B. H_2_O_2_ is the co-substrate for LPMO in driving its peroxygenase reaction, however, LPMOs are prone to autocatalytic oxidative inactivation in H_2_O_2_-driven reactions due to the generation of powerful oxygen species that will oxidatively damage the residues close to the catalytic copper (40–42). H_2_O_2_ could be generated not only by AA12 enzymes, but also by the oxidase activity of LPMOs. So, CAT was added as a H_2_O_2_ scavenger to eliminate the influence of H_2_O_2_ in this study. We found that the oxidized products of *Nc*LPMO9C were significantly increased as CAT added, further demonstrating H_2_O_2_-independent driving function of single-domain AA12 enzymes. It should be noting that the molecular size of free AA8 module is much smaller than the multi-domain CHDs or *Cc*PDH, so it could be easier to diffuse in the extracellular enzyme system during lignocellulose biodegradation process. So, we speculate that it may be more advantageous as an electron shuttle priming distant LPMOs, further work with different kinds of LPMOs will be needed to explore the interplay between LPMOs and free AA8 module in polysaccharide biodegradation (11,43).

## CONCLUSION

The single-domain AA12 PQQ-dependent oxidoreductases and free AA8 modules are prevalent in fungi. With aim to understand the interactions and functions of single-domain AA12 enzymes and free AA8 modules in fueling LPMO activity, three single-domain AA12 enzymes and one free AA8 modules were heterologously expressed and characterized in *P. pastoris*. All three tested single-domain AA12 enzymes are restrict dehydrogenases with trace residual oxidase activity. This study demonstrated that all three tested single-domain AA12 enzymes could directly transfer the electrons to Cu(II) in LPMO and drive *Nc*LPMO9C activity. The inter-protein electron transfer between single-domain AA12 enzymes and AA8 modules occurred, despite the electron transfer rate is lower than the inter-domain electron transfer in multi-domain CDHs. The AA12 enzyme-driven *Nc*LPMO9C efficiency could be significantly enhanced *by* the addition of free AA8 module *Tth*AA8B, probably attributing to the acceleration of electron transfer from AA12 enzymes to *Nc*LPMO9C and the attenuation of H_2_O_2_ accumulation mediated by *Tth*AA8B. Our findings highlight the critical role of single-domain AA12 enzymes and free AA8 modules in the biodegradation system of LPMOs.

## ACKNOWLEDGMENTS

This work was supported by the National Natural Science Foundation of China (NSFC) (22178179). The authors declare no competing financial interest.

AUTHOR CONTRIBUTIONS

Yuxing Sun: Investigation, design of experimental work, data visualization and analysis, writing original draft and editing. Congcong Yuan: Investigation. Liangkun Long: methodological guidance and validation. Shaojun Ding: conceptualization, supervision, funding acquisition, and writing review and editing. All authors read and approved the manuscript.

## ADDITIONAL FILES

The following material is available online.

**Supplemental Material** Figure S1-S8; Table S1-S3.

## ABBREVIATIONS

AA: auxiliary activity
AscA: Ascorbic acid
BMGY: buffered glycerol?complex medium
BMMY: buffered methanol?complex medium
CAT: catalase
CDH: cellobiose dehydrogenase
2,6-DMP: 2,6-dimethoxyphenol
EPR: electron paramagnetic resonance
FAD: flavin-adenine dinucleotide
H2O2: Hydrogen peroxide
HPAEC-PAD: high performance anion exchange chromatography-pulsed amperometric detection
HRP: horseradish peroxidase
LPMO: lytic polysaccharide monooxygenase
PASC: phosphoric acid-swollen cellulose
PDH: pyranose dehydrogenase
PQQ: pyrroloquinoline quinone
SDS-PAGE: SDS-polyacrylamide gel electrophoresis
YPD: yeast peptone dextrose medium.

